## Supplementary Figures S1-7 for "Single-cell RNA sequencing of *Plasmodium* vivax sporozoites reveals stage- and species-specific transcriptomic signatures"

* Corresponding author

**Supplementary Figures S1-7**

Fig S1

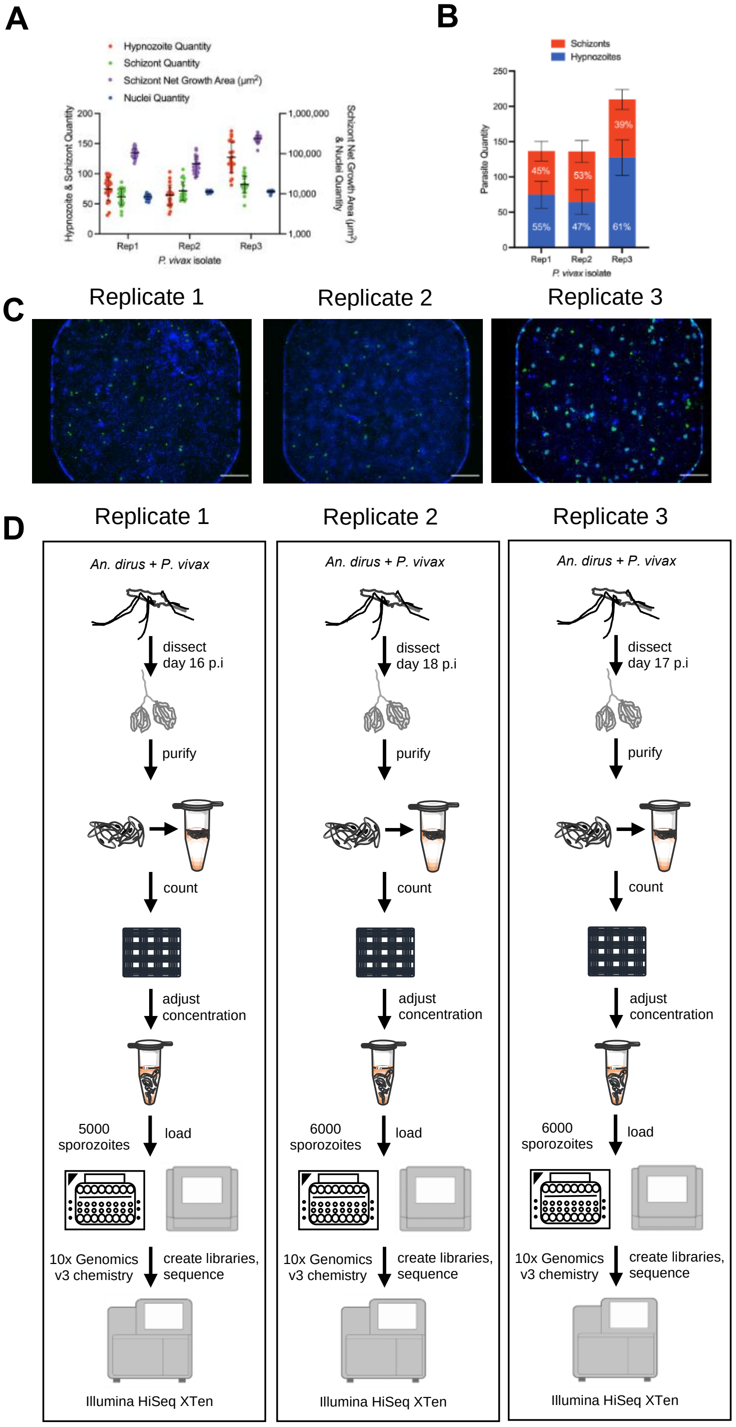


**Supplementary Figure 1. Related to Figure 1. Strategy used to assess *P. vivax* sporozoites transcriptomes at single-cell resolution.**
(A) Quantity of all hypnozoites, schizonts, and hepatic nuclei, as well as net growth area of schizonts, following infection of primary human hepatocytes with sporozoites from the indicated *P. vivax* replicate. Cultures were quantified at 12 days post infection. Each data point represents a single well of 24 technical replicate wells of a 384-well microtiter plate, bars represent SD across wells. The quantity of sporozoites infected into each well was 17,000 for Rep1, 16,500 for Rep2, and 18,000 for Rep3. (B) Ratio of hypnozoites versus schizonts for each isolate from ‘A,’ bars represent SD across 24 technical replicate wells. (C) Image of an individual culture well infected with sporozoites from the indicated isolate at 12 days post infection. Images are stitched from four fields of view taken at low-magnification (4x objective) during high-content imaging. Blue: Hoechst-stained host cell and parasite DNA, green: parasitophorous vacuole membrane detected with immunofluorescent staining with recombinant mouse anti-PvUIS4 antibody. Bar represents 500 μm. **(D)** Detailed schematic of the workflow for generating *P. vivax* single-cell RNA sequencing libraries. *P. vivax* sporozoites were manually dissected and purified by isolated the salivary glands of infected *An. dirus* mosquitoes. Sporozoites were harvested from three independent infections and three different days post-infectious blood-meal. scRNA-seq libraries were generated for the sporozoites using the 10x Genomics’ 3’ gene expression User Guide.

Fig S2

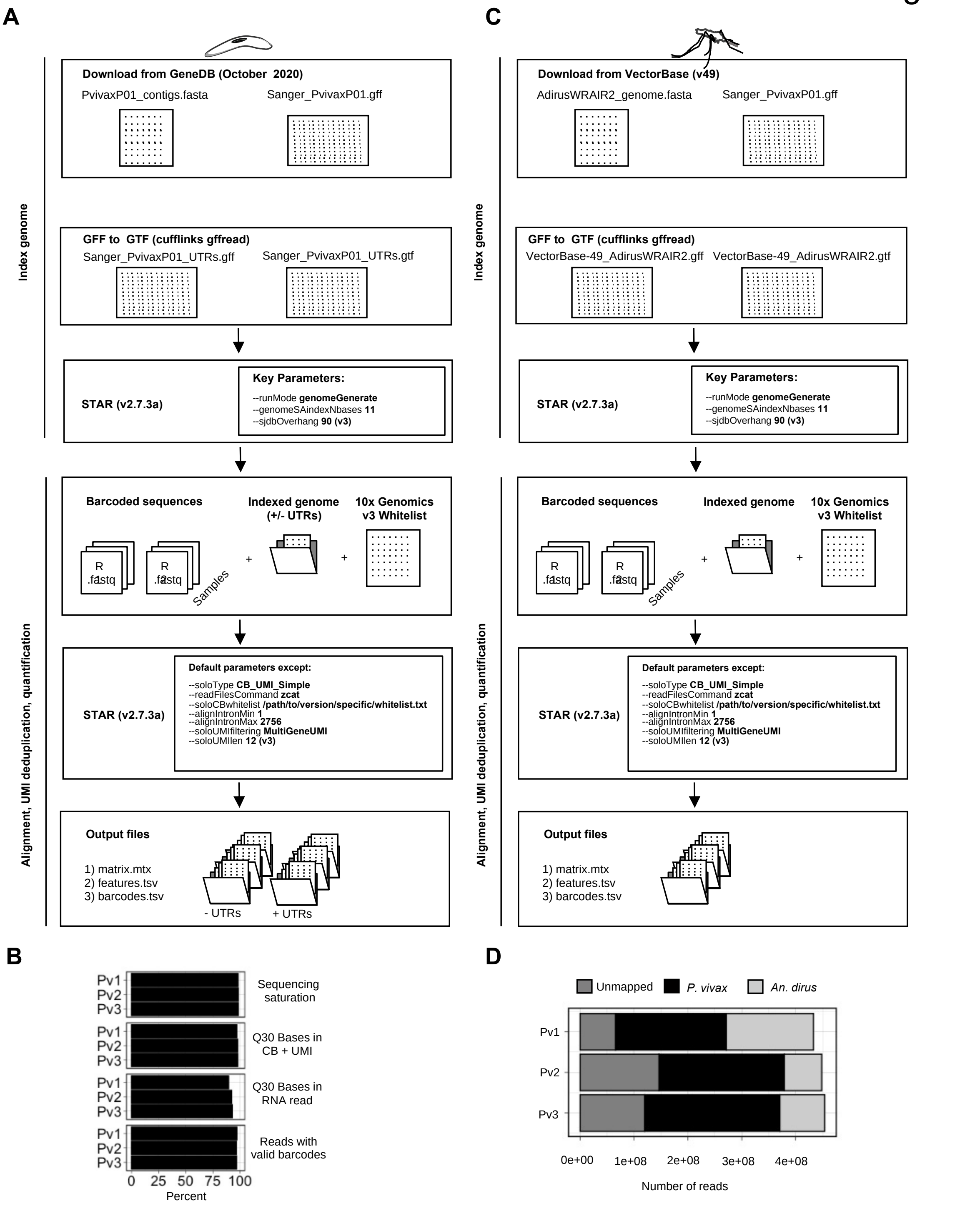


**Supplementary Figure 2. Related to Figure 1. Strategy used to assess *P. vivax* sporozoites transcriptomes at single-cell resolution.**
(A) Alignment of reads to *P. vivax* P01 genome, with parameters listed and output files at each stage. (B) Summary of output metrics from Illumina sequencing. (C) Alignment of reads to *An. dirus* WRAIR2 genome, with parameters listed and output files at each stage. (D) Number of reads mapping to *P. vivax* P01 genome, *An. dirus* WRAIR2 genome or unmapped to either.

Fig S3

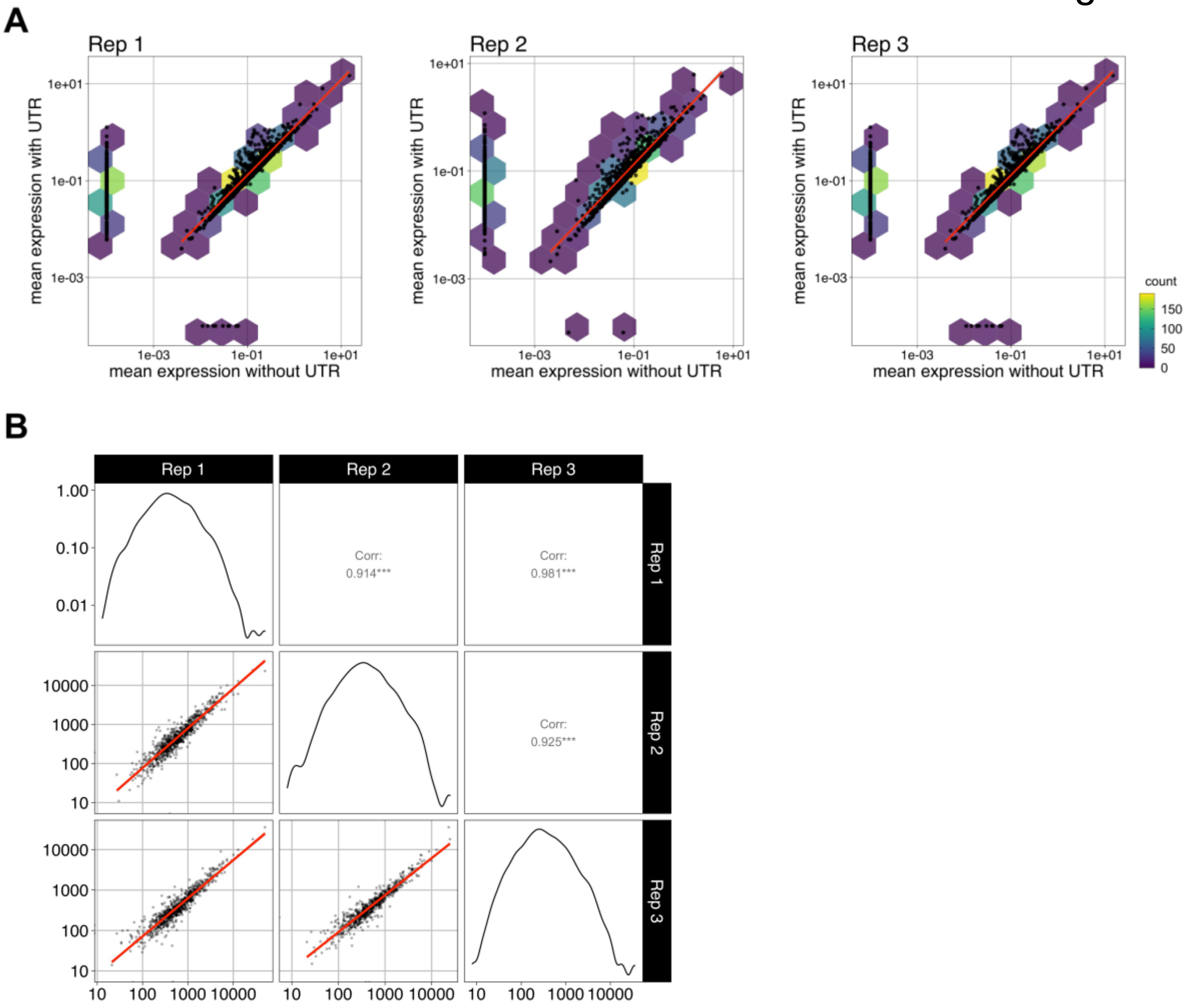


**Supplementary Figure 3. Related to Figure 2. Analysis of *P. vivax* sporozoite gene expression at single-cell resolution.** (A) Pairwise comparisons of transcript abundance (mean expression) with- or without- UTRs in gene models. (B) Pairwise comparisons of transcript abundance (mean expression) across the three replicates when sequencing reads are aligned to the *P. vivax* P01 genome with UTRs. Pearson's correlation coefficients (Corr, *R)* were determined using values > 0 for each pairwise comparison. ***, *p* value < 0.001.

Fig S4

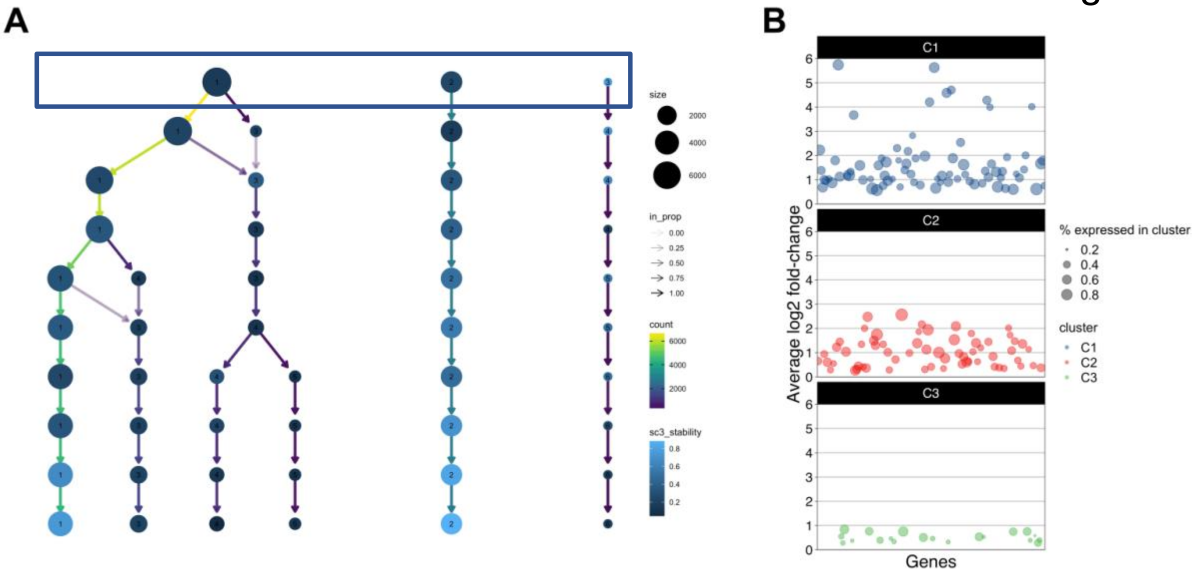


**Supplementary Figure 4. Related to Figure 2. Clustering and differential expression analysis of *P. vivax* sporozoites.** (A) Visualisation of cluster stability when resolution is increased in increments of 0.1 (start = 0.1 and end = 1.0).The size of each point is representative of the number of cells. Edges coloured by number of cells; and transparency represents the incoming node proportion (the number of samples in the edge divided by the number of samples in the node it points to). Point fill (sc3_stability) represents the calculated cluster stability. Clustering tree created with Clustree package (Zappia & Oshlack, 2018). Box indicates clustering resolution used for subsequent differential expression analysis. (B) Scatter plot of 159 genes identified as differentially expressed across the three clusters, split by the cluster. Genes displaying greater expression in each respective cluster plotted. Size of the point represents the percentage of cells expressing the gene of interest.

Fig S5

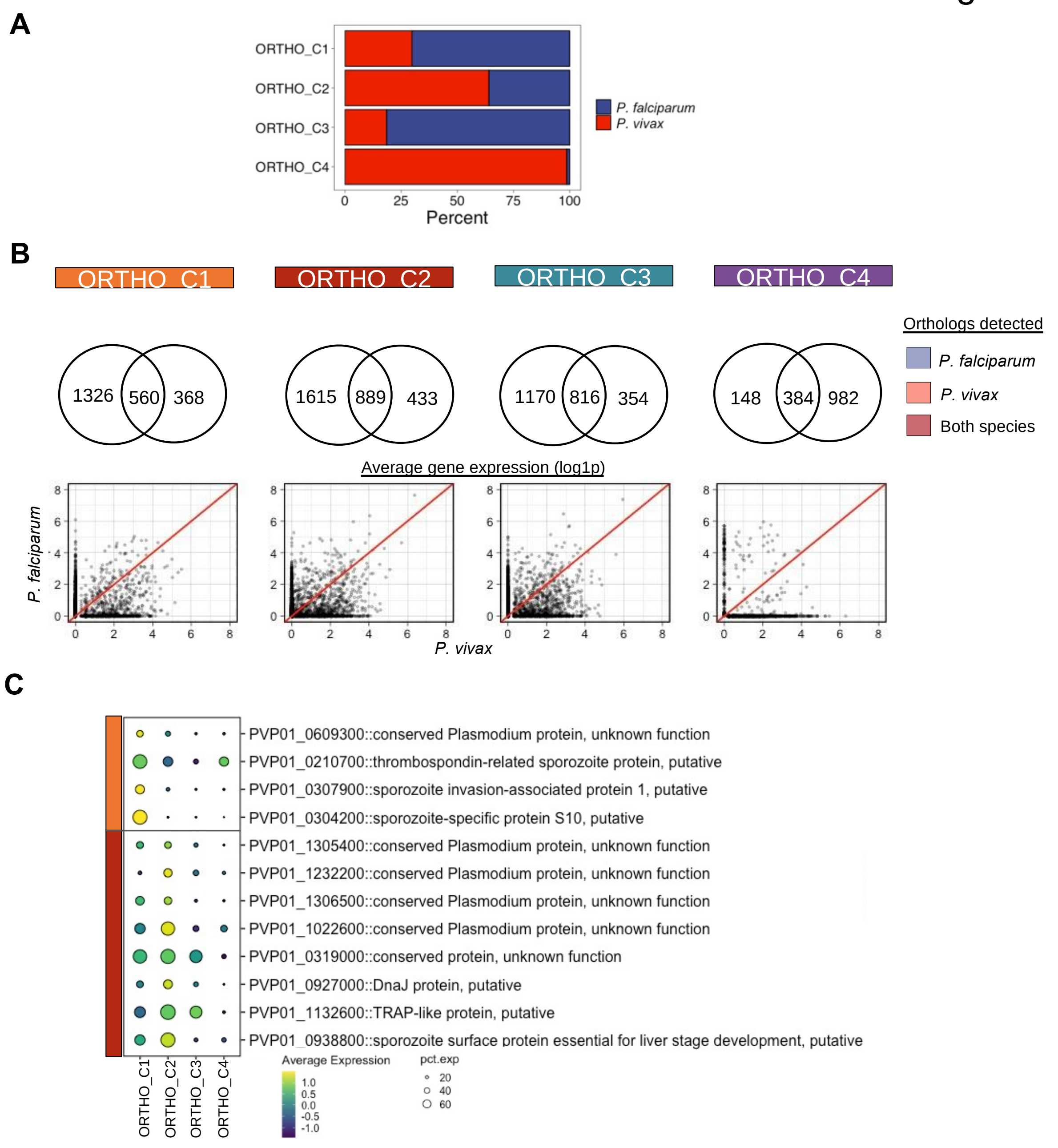


**Figure S5. Related to Figure 5. Integration of *P. vivax* and *P. falciparum* sporozoite datasets.**
(A) Proportion of cells derived from *P. vivax* and *P. falciparum* in each cluster. (B) Number of overlapping and unique one-to-one orthologs detected in each cluster. (C) Averaged gene expression (log1p) for genes detected in each cluster. (D) Dotplot of the top conserved genes across the two species for clusters one and two (Seurat parameters: min.pct .25, min.diff.pct = 0.125, Wilcoxon rank-sum test). Scale: Normalized expression, scaled; dot size: percentage of cells the transcript is detected.

Fig S6

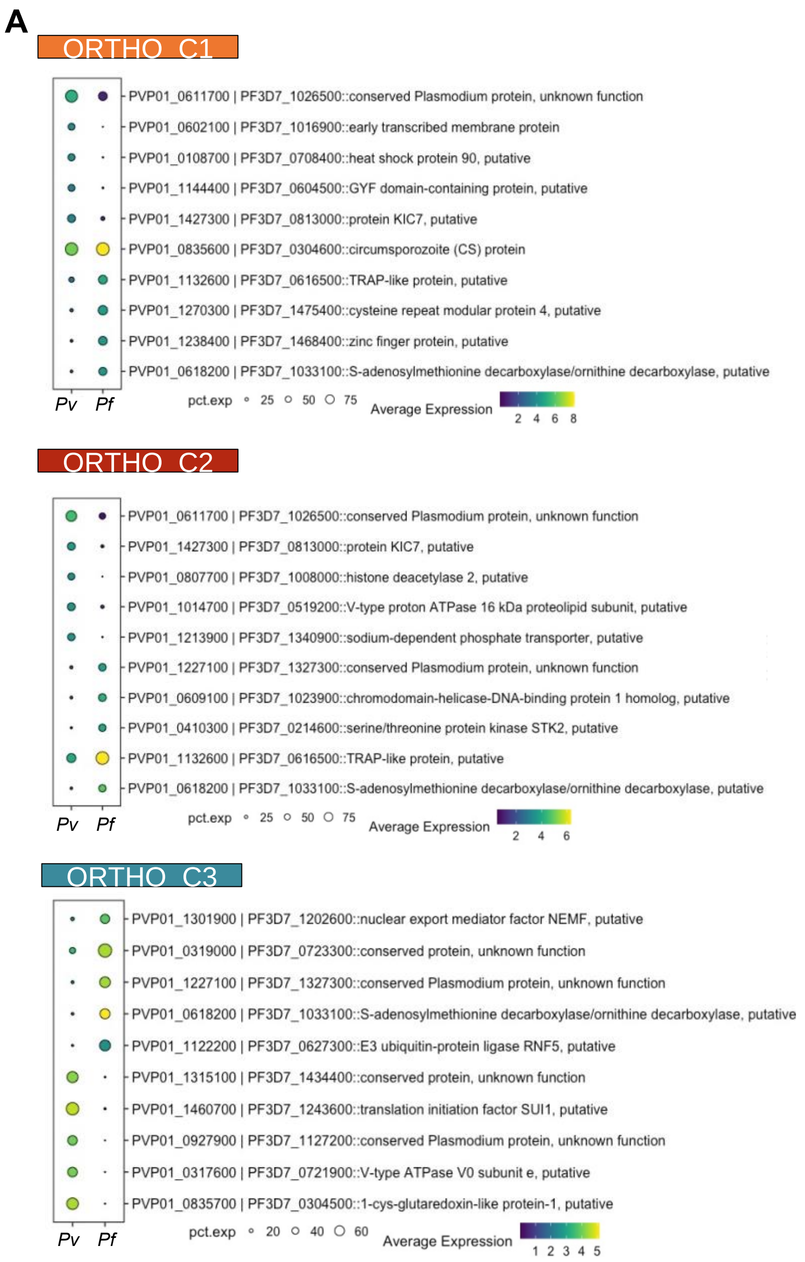


**Figure S6. Related to Figure 5. Integration of *P. vivax* and *P. falciparum* sporozoite datasets.**
(A) Dot plots of top one-to-one orthologs in each cluster that are differentially expressed between *P. vivax* and *P. falciparum.* The size of the dot corresponds to the percentage of cells expressing the gene. Scale bar: normalised expression.

Fig S7

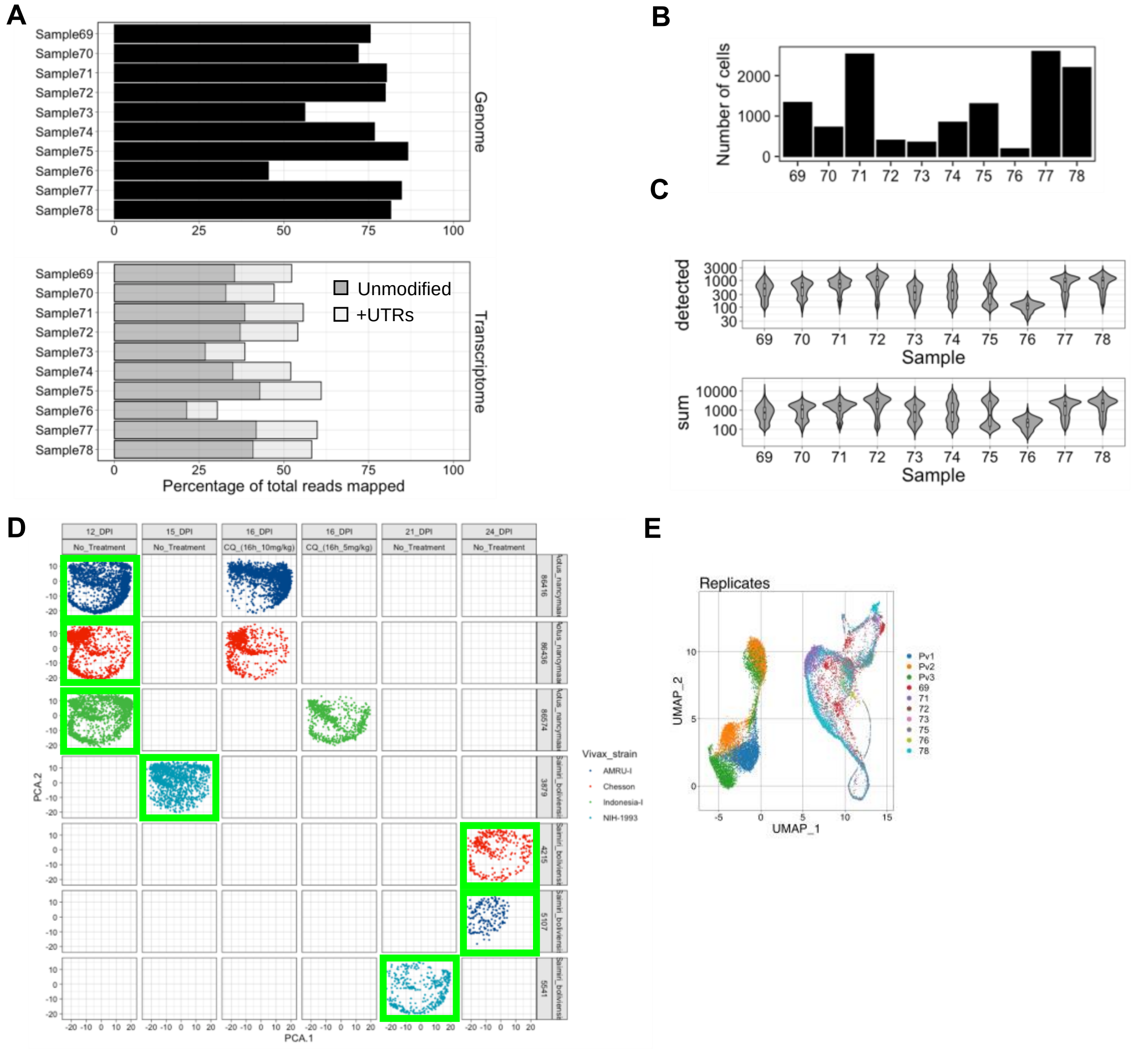


**Supplementary Figure 8. Related to Figure 6. Integration and comparative analyses of *P. vivax* sporozoite and blood-stage parasite transcriptomes.** Realignment, processing, and per-cell metrics of *P. vivax* blood stage 10x scRNAseq data prior to integration with the *P.vivax* sporozoite 10x scRNAseq data generated in this study.

(A) Percentage of reads aligning to the *P. vivax* P01 genome (upper panel) and transcriptome (with- or without- UTR information) (lower panel). (B) Number of *P. vivax* blood-stage transcriptomes retained post cell and gene filtering. (C) Violin plots showing the distribution of genes detected per cell (upper) and the UMIs detected per cell (lower). (D) PCA plots of the samples, split by day of infection, treatment, and monkey. Cells coloured by *P. vivax* strain used during monkey infection. Samples highlighted in green are those that are integrated with the scRNA-seq sporozoite data of the current study. (E) UMAP of integrated *P. vivax* sporozoite and blood stage data coloured by sample.
